# Retinoids rescue ceruloplasmin secretion and alleviate oxidative stress in Wilson’s disease-specific hepatocytes

**DOI:** 10.1101/2021.08.10.455792

**Authors:** Dan Song, Gou Takahashi, Yun-Wen Zheng, Mami Matsuo-Takasaki, Jingyue Li, Miho Takami, Yuri An, Yasuko Hemmi, Natsumi Miharada, Tsuyoshi Fujioka, Michiya Noguchi, Takashi Nakajima, Megumu K. Saito, Yukio Nakamura, Tatsuya Oda, Yuichiro Miyaoka, Yohei Hayashi

## Abstract

Wilson’s disease (WD) is a copper metabolic disorder, which is caused by defective ATP7B function. Here, we have generated induced pluripotent stem cells (iPSCs) from WD patients carrying compound heterozygous mutations on *ATP7B*. ATP7B loss- and gain-of-functions were further manifested with ATP7B-deficient iPSCs and heterozygously-corrected R778L WD patient-derived iPSCs using CRISPR-Cas9-based gene editing. Transcriptome analysis identified abnormalities of retinoid signaling pathway and lipid metabolism in WD-specific hepatocytes. Although the expression level of ATP7B protein was variable among WD-specific hepatocytes, the expression and secretion of ceruloplasmin (Cp), which is a downstream copper carrier in plasma, were consistently decreased. Cp secretion-based drug screening identified all-trans retinoic acid (ATRA) as promising candidates for rescuing Cp secretion. ATRA also alleviated reactive oxygen species (ROS) production induced by lipid accumulation in WD-specific hepatocytes. Our patient-derived iPSC-based hepatic models provide potential therapeutics for liver steatosis in WD and other fatty liver diseases.

## Introduction

Wilson’s disease (WD) (OMIM #277900), an autosomal recessive disorder, is majorly characterized by copper accumulation in the liver, leading to a series of metabolic disorders in the liver and nervous systems (Ala et al., 2007). WD is caused by defective *ATP7B*, which is located on chromosome 13, containing 21 exons, coding a protein of 1,465 amino acids known as copper-transporting P-type ATPase. More than 600 disease-causing mutations have been reported on ATP7B gene, which span almost all exons (Arioz et al., 2017; Huster et al., 2012), and also presents the pattern of specific mutations dominant in specific ethnic groups (Gomes and Dedoussis, 2016). The estimated incidence of WD is about 1:30000 (Ala et al., 2007; Gomes and Dedoussis, 2016), while the carrier frequency is up to 1 in 90. WD patients are facing lifelong medical treatments. Conventional therapies, such as penicillamine, trientine, zinc salt, and tetrathiomolybdate, caused severe side effects and significant variation in efficacy according to the cohort studies (Bandmann et al., 2015; Brewer, 2009; Huster et al., 2007; Kusuda et al., 2000). Thus, exploring new therapeutic medicine to prevent aggravation is urgent (Ferenci et al., 2003; Roberts et al., 2008).

To study the physiology and pathology of WD, immortalized cell lines (Polishchuk et al., 2014; van den Berghe et al., 2009) and rodent WD models (Buiakova et al., 1999; Rauch, 1983; Terada and Sugiyama, 1999; Yoshida et al., 1987) have been used conventionally; however, there is still a big gap between different species, as well as the different genetic backgrounds among individuals. Patient-derived iPSCs enable us to generate human hepatocytes to study WD physiology and pathology (Overeem et al., 2019; Parisi et al., 2018; Petters et al., 2020; Wei et al., 2020; Yi et al., 2012; Zhang et al., 2019; Zhang et al., 2020; Zhang et al., 2011). Previous studies majorly focused on high frequent hot spot mutations in WD patients (Coffey et al., 2013). However, due to limited numbers of patient numbers used in these studies, which fell short of statistical tests, generally targetable features of WD caused by variable mutations have not been fully identified.

In our present study, we generated iPSC lines from 4 patients carrying compound mutations on ATP7B gene. Also, we generated ATP7B-deficient iPSCs from a healthy-donor iPSC line by gene editing and mutation-corrected iPSCs from R778L homozygous mutant iPSCs generated previously (Overeem et al., 2019; Parisi et al., 2018; Yi et al., 2012; Zhang et al., 2011). With these WD-specific and ATP7B-deficient iPSC lines, we aimed to explore the disease phenotype-genotype correlation and to seek novel therapeutic candidates. We focused on ceruloplasmin (Cp), which is a copper-containing plasma ferroxidase synthesized in hepatocytes and secreted into plasma following incorporation of six atoms of copper by ATP7B protein in the trans-Golgi network (Sato and Gitlin, 1991). The low concentration of serum Cp is generally used as a diagnostic criterion of WD (Hellman and Gitlin, 2002; Saha et al., 2008). We successfully recapitulated the reduction of Cp expression and secretion in WD-iPSC-derived hepatocytes. From a small drug screening for rescuing Cp secretion levels in WD-iPSC-derived hepatocytes, we identified all-trans retinoic acid (ATRA) and clinically-approved retinoids, as promising candidates. Transcriptome analysis identified differentially-regulated genes in WD-specific hepatocytes that led to abnormalities of retinoid signaling pathways and lipid metabolism. Also, ATRA alleviated reactive oxygen species (ROS) production induced by lipid accumulation in oleic-acid-treated WD-specific hepatocytes.

Since Cp reduction and liver steatosis are the initial symptoms of WD, our results implied that retinoids could prevent or delay the progression of these symptoms.

## Results

### Generation and characterization of patient-derived WD-iPSCs

We generated patient-derived iPSC lines from 4 patients who suffered from WD, namely HPS0045, HPS0049, HPS0053, and HPS2807, by retroviral vectors or non-integrating episomal DNA vectors **(Table S1)**. All of these 4 WD-iPSC lines highly expressed self-renewal markers of iPSCs, such as OCT3/4, NANOG, SOX2, and KLF4 **(Figures 1A and S1A)**. Embryoid body (EB) formation assay, which allowed spontaneous differentiation of iPSCs into three germ layers *in vitro*, revealed pluripotency of 4 WD-iPSC lines **(Figure 1B)**. Also, teratomas containing various tissues derived from three germ layers were formed from these WD-iPSC lines **(Figure S1B)**. By comparative genome hybridization (CGH) array analysis, all of these WD-iPSC lines had normal karyotype without large copy number variations **(Figure 1C)**. As *ATP7B* was the responsible gene of WD (Frydman et al., 1985; Tanzi et al., 1993), we detected mutations in this gene carried in these WD-iPSC lines by sequencing the entire coding region of ATP7B from cDNA in WD-iPSC-derived hepatocyte cultures. We further verified the results with allelic identification partially in the cDNA and genomic DNA of these WD-iPSCs. Sequence results showed that HPS0045 iPSC line carried R778L (c.2333G>T), K832R (c.2495A>G) and R952K (c.2855G>A) heterozygous mutations. Both HPS0049 and HPS0053 iPSC lines carried G1035V (c.3104G>T) and V1262F (c.3784G>T) heterozygous mutations since they were siblings. HPS2807 iPSC line carried S406A (c.1216T>G), V456L (c.1366G>C), K832R (c.2495A>G), R952K (c.2855G>A), P992L (c.2975C>T) and K1010T (c.3029A>C) heterozygous mutations **(Figure 1D-1F)**. These characterization results indicated that these WD-iPSC lines maintained self-renewal, pluripotency, genomic integrity, and compound heterozygous mutations in *ATP7B* gene.

**Figure 1.**
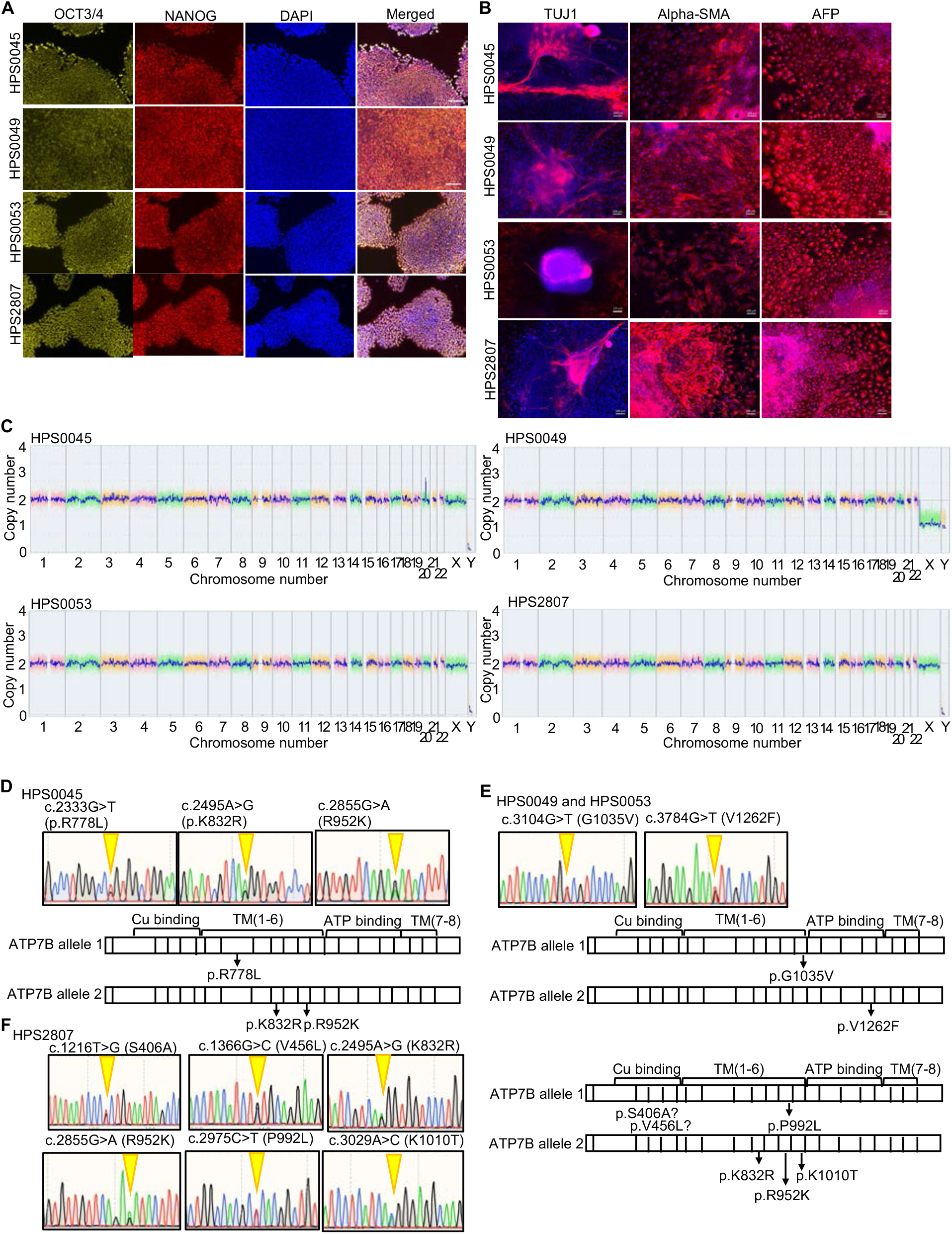
Generation of WD-patient-derived iPSC lines carrying compound heterozygous *ATP7B* mutations. (A) Expression of self-renewal markers of iPSCs, OCT3/4 (yellow) and NANOG (red), in WD-iPSCs. DAPI was used to stain nuclei (blue). Scale bar= 100 um. (B) Pluripotency was evaluated with EB formation assay. TUJ1 (ectoderm marker)/alpha-SMA (mesoderm marker)/AFP (endoderm marker) expression of EB. Scale bar=100 um. (C) Genome-wide copy number variation detection using CGH array analysis on WD-iPSC lines established in this study. (D-F) Sequence results and schema of allelic mutation positions of *ATP7B* genomic DNA in WD-patient-derived iPSC lines generated in this study.

### Hepatic differentiation of WD-iPSC lines

To establish disease models of WD *in vitro*, WD-iPSC lines and an iPSC line from a healthy donor (i.e., WTC11 line (GM25256)) were differentiated into hepatocytes with a monolayer differentiation protocol **(Figure S2A)**, which was modified from a protocol reported in a previous study (Siller et al., 2015). In our differentiation protocol, cells highly expressed definitive endodermal marker genes, *SOX17, HNF4A*, and *DLK1*, on days 4-7 **(Figure S2B)**, and hepatoblast and/or hepatocyte marker genes, alpha-fetoprotein (*AFP)*, albumin (*ALB)*, and alpha-1-antitrypsin (*A1AT)*, on day 17 **(Figure S2C)**. Immunostaining results showed that definitive endodermal marker proteins, SOX17 and GATA6, were highly expressed on day 5 **(Figures S2D and S2E)** and that hepatic marker proteins, HNF4A, AFP, and ALB were highly expressed on day 17 **(Figures S2F-S2H)**. WD-specific hepatocytes also expressed comparable levels of hepatic marker genes and proteins to WTC11-Hep (hepatocytes differentiated from WTC11 iPSC line) on differentiation day 17 **(Figures S3A-S3D)**. These results indicated the comparable differentiation ability between WD-specific iPSCs and WTC11 iPSCs into hepatic lineages and thus the suitability of our present hepatic differentiation protocol for WD modeling *in vitro*.

### CP expression is consistently decreased in WD-iPSC-derived hepatocytes

We asked how WD mutations affected ATP7B mRNA and protein expression in WD-iPSC-derived hepatocytes. In our differentiation protocol, *ATP7B* mRNA expression level constantly increased **(Figure S3E)**. Thus, the expression levels of ATP7B mRNA and protein were examined in WD-specific hepatocytes on differentiation day 17 by RT-qPCR and western blot, respectively. All the WD-specific hepatocytes expressed a comparable level of *ATP7B* mRNA with WTC11-derived hepatocytes **(Figure 2A)**. At the protein level, HPS2807-derived hepatocytes expressed a similar amount of ATP7B protein compared to WTC11-derived hepatocytes; HPS0045-derived hepatocytes expressed a lower (but not a significant) amount of ATP7B protein; HPS0049- and HPS0053-derived hepatocytes expressed almost no ATP7B protein **(Figures 2B and 2C)**. These results suggested that WD-specific iPSC-derived hepatocytes expressed highly variable levels of ATP7B protein regulated in the post-transcriptional processes depending on the types of ATP7B mutations.

**Figure 2.**
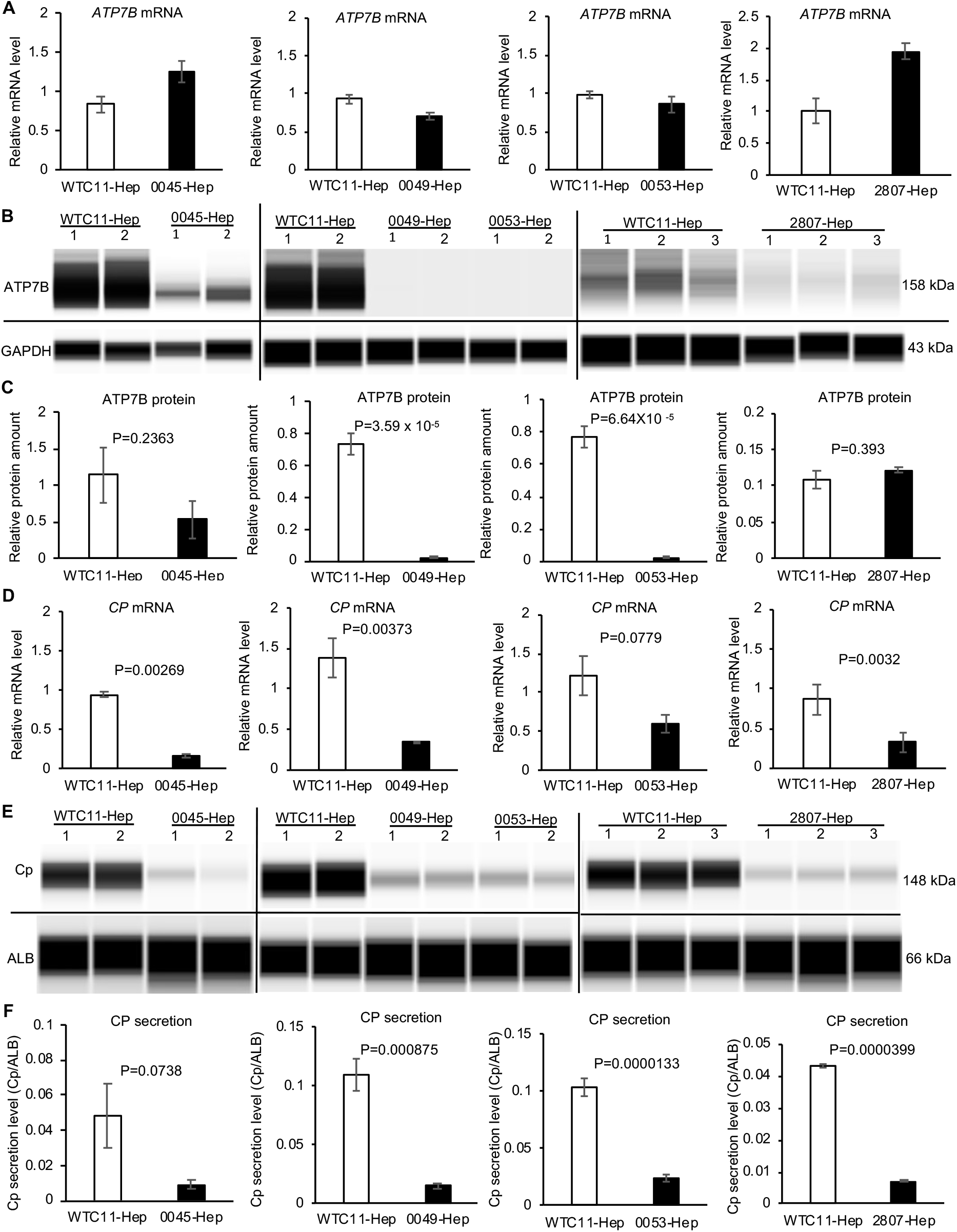
The levels of expression and secretion of Cp are consistently decreased in WD-iPSC-derived hepatocytes. (A) Expression levels of *ATP7B* mRNA in WD-iPSC-derived hepatocytes on differentiation day 17 quantified by RT-qPCR (0045-Hep, n=3; 0053-Hep, n=3; 0049-Hep, n=4; 2807-Hep, n=3). Data are shown as mean ± SEM (standard error of the mean). (B) Representative images of western blot results of ATP7B protein expression in WTC11- and WD-iPSC-derived hepatocytes (WTC11-Hep, 0045-Hep, 0049-Hep, and 0053-Hep, respectively) on differentiation day 17. GAPDH protein was used as a housekeeping control. (C) Bar graphs showing quantified expression levels of ATP7B protein normalized with the expression levels of GAPDH protein. The data obtained from WTC11-Hep were used as the common control for each WD-Hep (0045-Hep, n=4; 0053-Hep, n=6; 0049-Hep, n=7; 2807-Hep, n=3). Data are shown as mean ± SEM. P-values were determined by unpaired two-tailed Student’s *t*-test. (D) Expression levels of *CP* mRNA in WTC11-Hep and WD-iPSC-derived-Hep. Data are shown as mean ±SEM. P-values were determined by unpaired two-tailed Student’s *t*-test. (E) Representative images of western blot data of secreted Cp and ALB proteins by WTC11- and WD-iPSC-derived hepatocyte cultures on differentiation day 17. (F) Quantified data of Cp protein secretion levels after normalized with ALB. WTC11-Hep was used as the common control for each WD-Hep (0045-Hep, n=7; 0053-Hep, n=7; WD0049-Hep, n=6; 2807-Hep, n=3). Data are shown as mean ± SEM. P-values were determined by unpaired two-tailed Student’s *t*-test.

Next, we examined mRNA expression and secretion level of Cp, which received copper ions by ATP7B protein in hepatocytes and is secreted into the plasma to deliver these ions to the whole body. The expression level of Cp was considerably increased during differentiation into hepatocytes from undifferentiated iPSCs **(Figure S3F)**. In iPSC-derived hepatocytes on differentiation day 17, the expression level of Cp mRNA was generally lower in WD-specific hepatocytes compared with WTC11-derived hepatocytes **(Figure 2D)**. Also, the level of Cp secretion, which was normalized by the amount of secreted ALB to evaluate Cp secretion levels per hepatocyte, was generally lower in WD-specific hepatocytes compared with WTC11-derived hepatocytes **(Figures 2E and 2F)**.

These results suggested that WD mutations in *ATP7B* generally resulted in reduced mRNA expression and protein secretion of Cp in iPSC-derived hepatocytes.

### Gene-editing of *ATP7B* in iPSC-derived hepatocytes confirmed the regulation of Cp expression and secretion by ATP7B

To verify the effect of loss-of-function of *ATP7B* with isogenic backgrounds in iPSC-derived hepatocyte models, we generated *ATP7B*-deficient iPSC clones from WTC11 iPSC line using CRISPR-Cas9 system (Ran et al., 2013b). The translational start codon of *ATP7B* gene was targeted by a guide RNA (gRNA) **(Figure 3A)**. After introducing the gRNA and Cas9 into the WTC11 iPSC line, we have isolated iPSC clones, in which 7 bp was deleted in *ATP7B* exon 1 in one allele, and 104 bp spanning exon 1 and intron 1 was deleted in the other allele, detected by genotyping sequencing (**Figure 3B)**. The expression of hepatic markers expression was comparable between ATP7B-deficient and original WTC11-derived hepatocytes on differentiation day 17 **(Figures S4A-S4G)**. ATP7B gene and protein expression was examined in these ATP7B-deficient hepatocytes with RT-qPCR and western blotting, respectively. These ATP7B-deficient hepatocytes showed comparable ATP7B gene and protein expression levels as WTC11-derived hepatocytes **(Figures 3D and 3E)**. These results indicated that gene editing in *ATP7B* exon 1 did not interfere ATP7B expression in both transcriptional and post-transcriptional levels, possibly due to the generation of alternative start codons after its gene edition. Nonetheless, mRNA expression level and secretion level of Cp were significantly lower in ATP7B-deficient hepatocytes **(Figures 3F and 3G)**. These results indicated that the functionality of the ATP7B protein expressed in these ATP7B-deficient hepatocytes was severely compromised.

**Figure 3.**
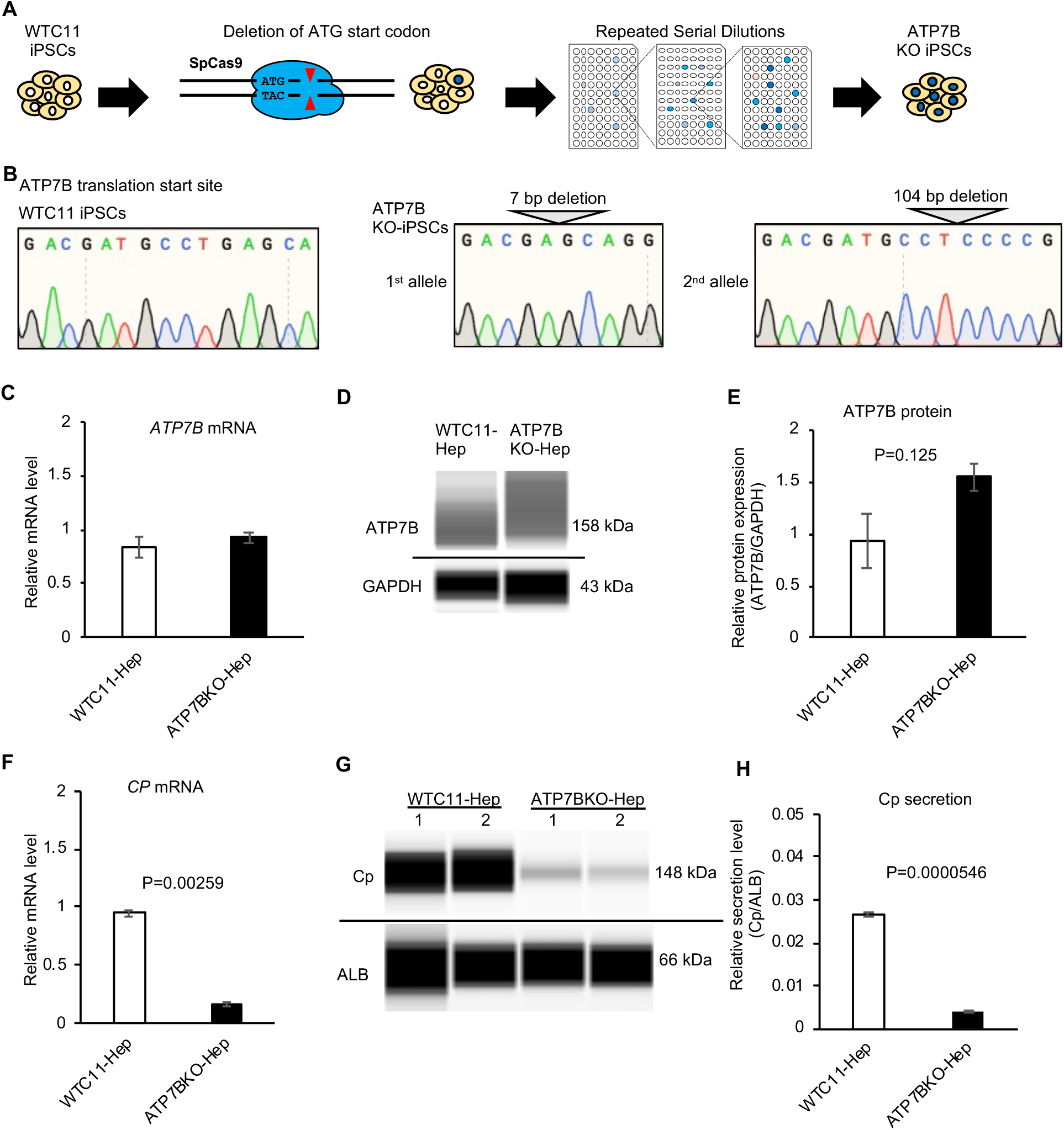
The levels of expression and secretion of Ceruloplasmin are decreased in *ATP7B*-deficient-iPSC-derived hepatocytes. (A) Schema of the generation of *ATP7B*-deficient iPSCs by gene-editing technology. (B) Allele-specific sequence data around the start codon of *ATP7B* genomic DNA in a gene-edited iPSC clone. (C) The expression level of *ATP7B* mRNA in WTC11-Hep and ATP7B-deficient-Hep. Data are shown as mean ± SEM (n=3). (D) Representative image of western blot data of ATP7B and GAPDH proteins from parental WTC11- and ATP7B-deficient-iPSC-derived hepatocyte cultures on differentiation day 17. (E) Quantified data of ATP7B protein expression levels after normalized with GAPDH protein. Data are shown as mean ± SEM (n=3). (F) The expression level of *CP* mRNA in WTC11-Hep and ATP7B-deficient-Hep. Data are shown as mean ± SEM (n=3). P-values were determined by unpaired two-tailed Student’s *t*-test. (G) Representative image of western blot data of secreted Cp and ALB proteins by WTC11- and ATP7B-deficient-Hep on differentiation day 17. (H) Quantified data of secreted CP protein levels after normalized with ALB protein in WTC11-Hep and ATP7B-deficient-Hep. Data are shown as mean ± SEM (n=3). P-values were determined by unpaired two-tailed Student’s *t*-test.

Conversely, to examine the effect of genetic correction of ATP7B with isogenic backgrounds on WD disease modeling, we performed CRISPR-Cas9-based gene correction on one WD-iPSC line carrying predominated homozygous mutation (R778L), which was generated previously(Zhang et al., 2011) **(Figure 4A)**. Near R778L mutation site was targeted by a gRNA with Cas9 D10A nickase, and an oligonucleotide DNA was used as a homologous recombination donor to generate scarlessly rescued iPSC subclones (Ran et al., 2013a). After introducing this gRNA, ssODN, and Cas9 D10A nickase into R778L-homozygous iPSCs, we have isolated single-cell clones and selected R778L-heterozygous iPSC subclones. The genomic DNA sequence of the exon in ATP7B gene was confirmed by Sanger sequencing (**Figure 4B)**. Hepatic differentiation capability of the corrected R778L-heterozygous iPSC subclone and the original R778L-homozygous iPSC line was examined by immunostaining and RT-qPCR. These two iPSC subclones expressed comparable levels of hepatic markers **(Figures S5A-S5F)**. ATP7B mRNA and protein levels were examined in these iPSC-derived hepatocytes. While both iPSC clones expressed similar levels of *ATP7B* mRNA, this R778L-heterozygous line expressed a significantly higher level of ATP7B protein **(Figures 4C-4E)**. Cp mRNA and secretion levels were further examined in these conditions. This R778L-heterozygous subclone expressed significantly higher levels of mRNA and secreted protein of Cp than those of the original R778L-homozygous line **(Figures 4F-4H)**. These results suggested that heterozygous correction of R778L rescued Cp expression and secretion levels as well as ATP7B protein expression and function. Collectively, our results demonstrated that loss- and gain-of-function of ATP7B manifested the regulation of Cp expression and secretion levels in iPSC-derived hepatocytes.

**Figure 4.**
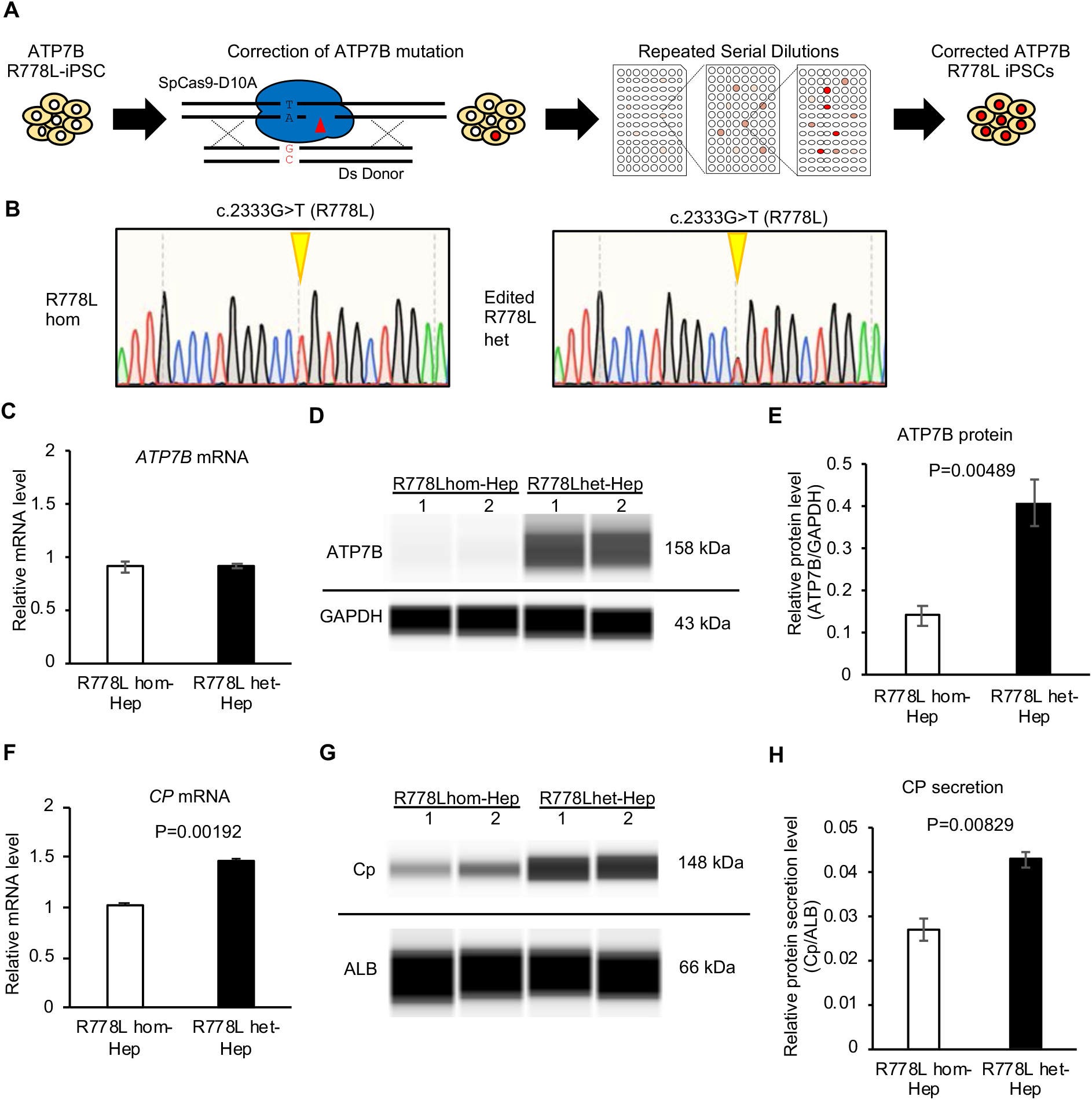
The levels of expression and secretion of Ceruloplasmin are increased in heterozygously-corrected ATP7B R778L-iPSC-derived hepatocytes. (A) Schema of the generation of heterozygously-corrected *ATP7B* R778L iPSCs by gene-editing technology. (B) Sequence results around the R778L position of *ATP7B* genomic DNA in parental R778L homozygous iPSCs (R778Lhom) and the established gene-edited iPSC clone (R778Lhet). (C) The expression level of *ATP7B* mRNA in WTC11-Hep and ATP7B-edited-Hep on differentiation day 17. Data are shown as mean ± SEM (n=3). (D) Representative image of western blot data of ATP7B and GAPDH proteins from parental R778Lhom- and R778Lhet-iPSC-derived hepatocytes on differentiation day 17. (E) Quantified data of ATP7B protein expression levels after normalized with GAPDH protein. Data are shown as mean ± SEM (n=3). (F) Expression levels of *CP* mRNA in R778Lhom-Hep and R778Lhet-Hep detected by RT-qPCR. Data are shown as mean ± SEM (n=3). P-values were determined by unpaired two-tailed Student’s *t*-test. (G) Representative image of western blot data of secreted Cp and ALB proteins by R778Lhom-Hep and R778Lhet-Hep on differentiation day 17. (H) Quantified data of secreted CP protein levels after normalized with ALB protein in R778L hom-Hep and R778L het-Hep. Data are shown as mean ± SEM (n=3). P-values were determined by unpaired two-tailed Student’s *t*-test.

### Retinoids were identified to increase Cp expression and secretion levels in WD-specific hepatocytes

From our results above, the decreased level of secreted Cp is a robust and convenient indicator of ATP7B dysfunction in WD-specific iPSC-derived hepatocytes. Because up-regulating Cp secretion capability of these WD-specific hepatocytes should be helpful to relieve excessive copper-induced symptoms, we developed a chemical screening system to evaluate Cp secretion levels in iPSC-derived hepatocytes using an automatic capillary western blot device **(Figure 5A)**. In this study, eleven candidate chemicals, which had been reported to affect Cp expression and/or secretion in other biological contexts, were screened in these five WD-specific iPSC-derived hepatocytes. Among these 11 candidate chemicals, only ATRA significantly increased Cp secretion levels of these hepatocytes in the comparison with DMSO-treated control conditions **(Figures 5B, 5C, and S6A)**. The addition of ATRA increased *CP* mRNA expression without obvious changes in the expression of other hepatic genes in these 5 WD-specific hepatocytes with RT-qPCR **(Figure 5D)**. These results suggested that ATRA rescued decreased expression and secretion of Cp in WD-specific hepatocytes without affecting hepatocyte identity.

**Figure 5.**
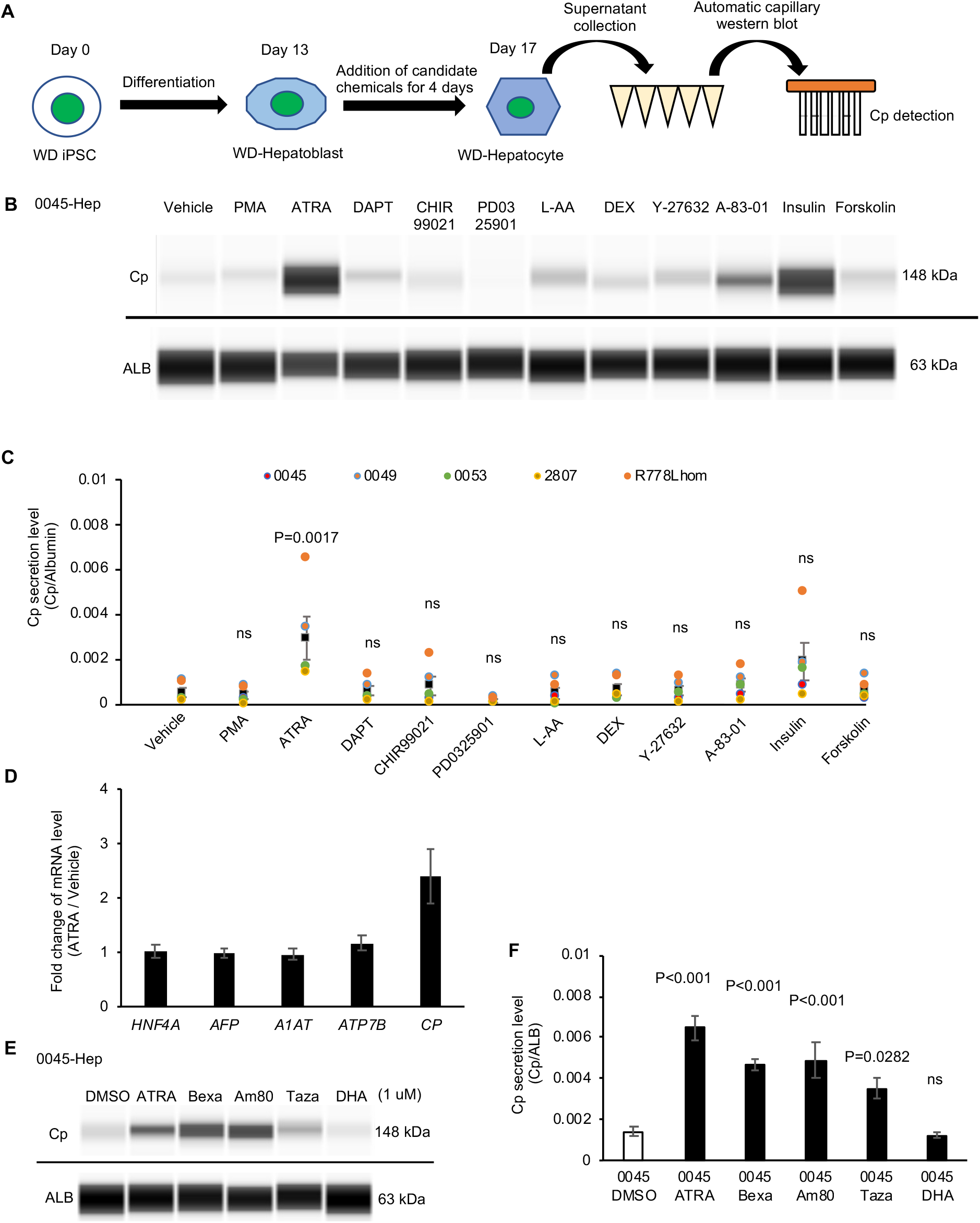
Drug screening identifies ATRA and retinoids to increase CP expression and secretion in WD-iPSC-derived hepatocytes. (A) Schema of Cp secretion-based drug screening. (B) Representative image of western blot data of secreted Cp and ALB proteins by 0045-Hep on differentiation day 17 after treated with indicated drugs for 4 days. (C) Quantified data of Cp secretion level after normalized with ALB, dot colors indicate data from different WD-Hep. Data are shown as mean ± SEM (n=5). Statistical significance was determined by Dunnett’s test relative to DMSO-treated conditions (vehicle). (D) Expression levels of *HNF4A, AFP, A1AT, ATP7B, and CP* mRNA in 0045-Hep on differentiation day 17 after treated with DMSO (vehicle) or ATRA for 4 days quantified by RT-qPCR. Data are shown as mean ± SEM. (E) Representative images of western blots and quantification data showing Cp secretion level of 0045-Hep on day 17 after being treated with indicated two RXRs and two RARs agonists for 4 days. (F) Quantified data of secreted CP protein levels after normalized with ALB protein in 0045-Hep treated with indicated two RXRs and two RARs agonists. Data are shown as mean ± SEM (n=4). P-values were determined by Dunnett’s test relative to DMSO-treated condition (vehicle).

ATRA belongs to retinoid family molecules and regulates a wide variety of physiological functions through its nuclear receptors, known as the classical retinoic acid receptors (RARs) and nonclassical retinoid X receptors (RXRs) (Duester et al., 2003; Mangelsdorf et al., 1990). To further verify and extend the effect of ATRA on secret levels of Cp in WD-Hep clinically-used retinoids, we employed 4 clinically-approved retinoid derivatives; Tazarotene and Am80 are RAR-selective retinoids (Chandraratna, 1996; Nkongolo et al., 2019), and Bexarotene and cis-4,7,10,13,16,19-Docosahexaenoic acid (DHA) are RXR-selective retinoids (Mounier et al., 2015; Zapata-Gonzalez et al., 2008). Effects of these four retinoids and ATRA on Cp secretion were further examined in WD-specific hepatocytes. These retinoids except for DHA increased Cp secretion in hepatocytes derived from iPSC lines of HPS0045, HPS0053, and R778L-homozygous mutant **(Figures 5E, 5F, and S6B-S6E)**. These results suggested that retinoid derivatives rescued decreased Cp expression and secretion in WD-specific hepatocytes.

### Transcriptome analysis identified abnormalities of RA signaling pathway and lipid metabolism in WD-specific hepatocytes

To identify the differences in global gene expression patterns in WD-specific hepatocytes, we performed RNA-seq analysis. Six WD-specific hepatocytes and 6 healthy-donor (HD) hepatocytes were compared in total **(Figure 6A)**. From this comparison, 238 genes were significantly down-regulated (FDR < 0.05), and 431 genes were significantly up-regulated (FDR < 0.05) **(Figure 6B and Table S2)**. Of note, CP was included in these significantly down-regulated genes. Then, we performed enrichment analysis of biological pathways and gene ontology on these differentially-regulated genes. From these down-regulated genes, RA was the top metabolite enriched in these genes identified in the HMDB (Human Metabolite DataBase) program **(Figure 6C)**. From these up-regulated genes, lipid and lipoprotein metabolisms were the top pathways enriched in these genes identified in NCATS (National Center for Advancing Translational Sciences) BioPlanet program **(Figure 6D)**. Also, liver neoplasms, fatty liver, and steatohepatitis were the top associated diseases enriched in these genes identified in DisGeNET program **(Figure 6E)**. To examine these differentially-regulated genes in ATP7B deficiency contexts, we performed an integrated analysis of these genes with previous data sets from the samples of mouse liver comparing ATP7B knockout mice with wild-type mice (n=4). We counted overlapped genes that were significantly down-regulated and up-regulated in our WD-specific hepatocytes with a dataset from ATP7B knockout mouse liver samples (n=4) (i.e., GSE125637) (Wooton-Kee et al., 2020). Eighteen down-regulated genes out of 238 genes (7.6%) were overlapped with this dataset from mouse liver samples **(Figure 6F)**. Overlapped down-regulated genes included RXRG, TNNT2, FITM1, CCDC181, GPM6A, PSRC1, CKS2, TTK, BIRC5, DLGAP5, KIF2C, CDK1, CCNA2, UBE2C, HMGB2, CDC20, PLK1 and PRC1. Fifty up-regulated genes out of 431 genes (11.6%) were overlapped with this dataset from mouse liver samples **(Figure 6G)**. Overlapped up-regulated genes included AFF1, ZHX3, ALDH3A2, SOD2, GBF1, INSR, TXNDC11, OGFRL1, ELL2, BDH1, TSKU, GGCX, EGFR, ST6GAL1, CPEB4, HNF4A, MERTK, SLC5A9, ACOX1, TRIM14, PPARA, PPM1L, IL1RAP, CHPT1, PLXNC1, MICAL2, AR, DEPTOR, LIPC, S100A1, AK4, ABCA1, AQP3, ABCC3, KCNJ16, ACADL, SLC7A4, CLDN1, A1CF, RBP4, APOB, SERPINA7, SLC34A2, KLB, SLC10A1, ZFYVE28, C6, ABCB11, HOXC10, and HOXA9. These results suggested that our RNA-seq analysis was partially validated in the context of ATP7B deficiency and that these overlapped genes might be regulated by ATP7B deficiency among mammalian species in common.

**Figure 6.**
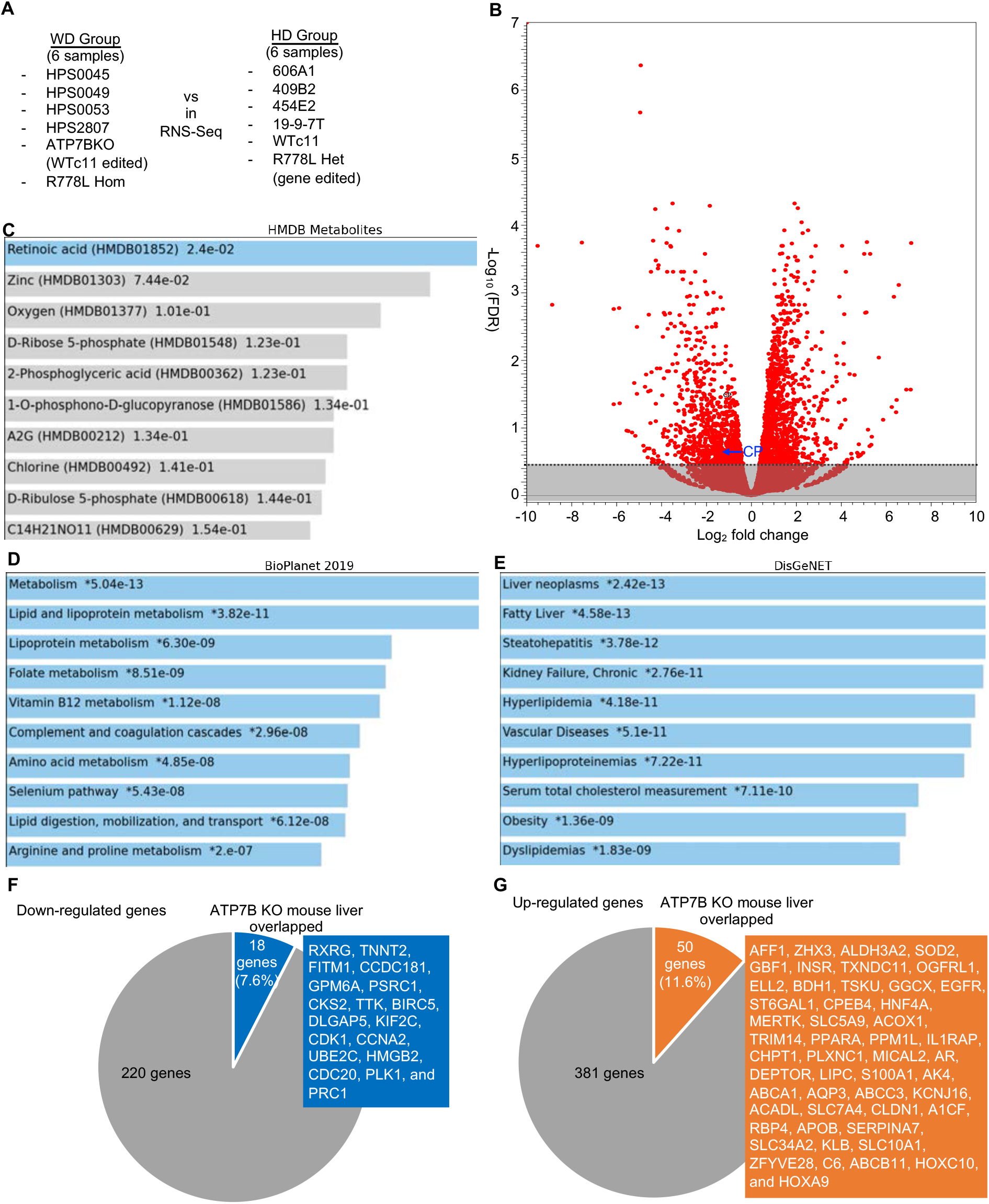
Transcriptome analysis identifies abnormality of RA signal pathway and lipid metabolism in WD-iPSC-derived hepatocytes. (A) Schema of RNA-seq. (B) Volcano plot shows Log2 fold change as X-axis and -log10 (FDR) as Y-axis. All the genes were plotted in red circles. CP gene was indicated by a blue arrow. Non-significant genes with FDR >= 0.05 were masked in gray. (C) A bar chart showing candidate metabolites using HMDB (human metabolome database) Metabolites program from down-regulated genes. Metabolite names (HMDB metabolite ID) and values of -log_10_(P-value) are shown. (D) A bar chart showing candidate pathways using NCATS (National Center for Advancing Translational Sciences) BioPlanet program from up-regulated genes. Pathway names and values of -log_10_(P-value) are shown. (E) A bar chart showing candidate disease types using DisGeNET program from up-regulated genes. Disease names and values of -log_10_(P-value) are shown. (F) A pie chart showing overlapped down-regulated genes in WD-specific hepatocytes with ATP7B KO (knockout) mouse liver samples. The number of these genes, the percentage of the overlapped genes, and the overlapped gene name are shown. (G) A pie chart showing overlapped up-regulated genes in WD-specific hepatocytes with ATP7B KO (knockout) mouse liver samples. The number of these genes, the percentage of the overlapped genes, and the overlapped gene name are shown.

### ATRA suppresses oleic acid-induced reactive oxygen species (ROS) production in WD-specific hepatocytes

Our transcriptome analysis on WD-specific hepatocytes above indicated that abnormal gene expression related to liver dysfunctions and lipid metabolism. Hepatic steatosis is one of the earliest symptoms in WD patients, which may be caused by abnormal lipid metabolism in response to copper accumulation in WD patients (Hamilton et al., 2016; Seessle et al., 2011; Wilkins et al., 2013). Although molecular mechanisms between copper accumulation and abnormal lipid metabolism in WD remain elusive, ROS (reactive oxygen species) plays a central role to mediate them (Chen et al., 2020; Hamilton et al., 2016; Seessle et al., 2011). Thus, to recapitulate features of hepatic steatosis in WD, lipid accumulation and ROS production were evaluated in WD-specific hepatocytes treated with different concentrations of oleic acid by fluorescent imaging with specific chemical probes (**Figure 7A**). In this assay system, we examined the effect of ATRA on lipid accumulation and ROS production. Lipid droplets were enlarged and outstanding in the WD-derived hepatocytes treated with higher concentrations of oleic acid **(Figures 7B and 7C)**. The level of ROS production was also higher in WD-specific hepatocytes treated with higher concentrations of oleic acid. Calculation of fluorescent intensity from these tiled whole-well images indicated that the treatment of ATRA reduced ROS production in the oleic acid-treated WD-derived hepatocytes, but not in the oleic acid-treated WTc11-derived hepatocytes **(Figures 7D-7F and S7A-S7C)**. Expression of hepatic marker genes, such as ALB, AFP, A1AT, and CP, was decreased by the addition of high concentration (200 µM) of oleic acid in WD-specific hepatocytes, but not in WTC11-derived hepatocytes **(Figure S7D and S7E)**. Our results indicated that ATRA alleviated ROS production induced by accumulated lipids in WD-derived hepatocytes.

**Figure 7.**
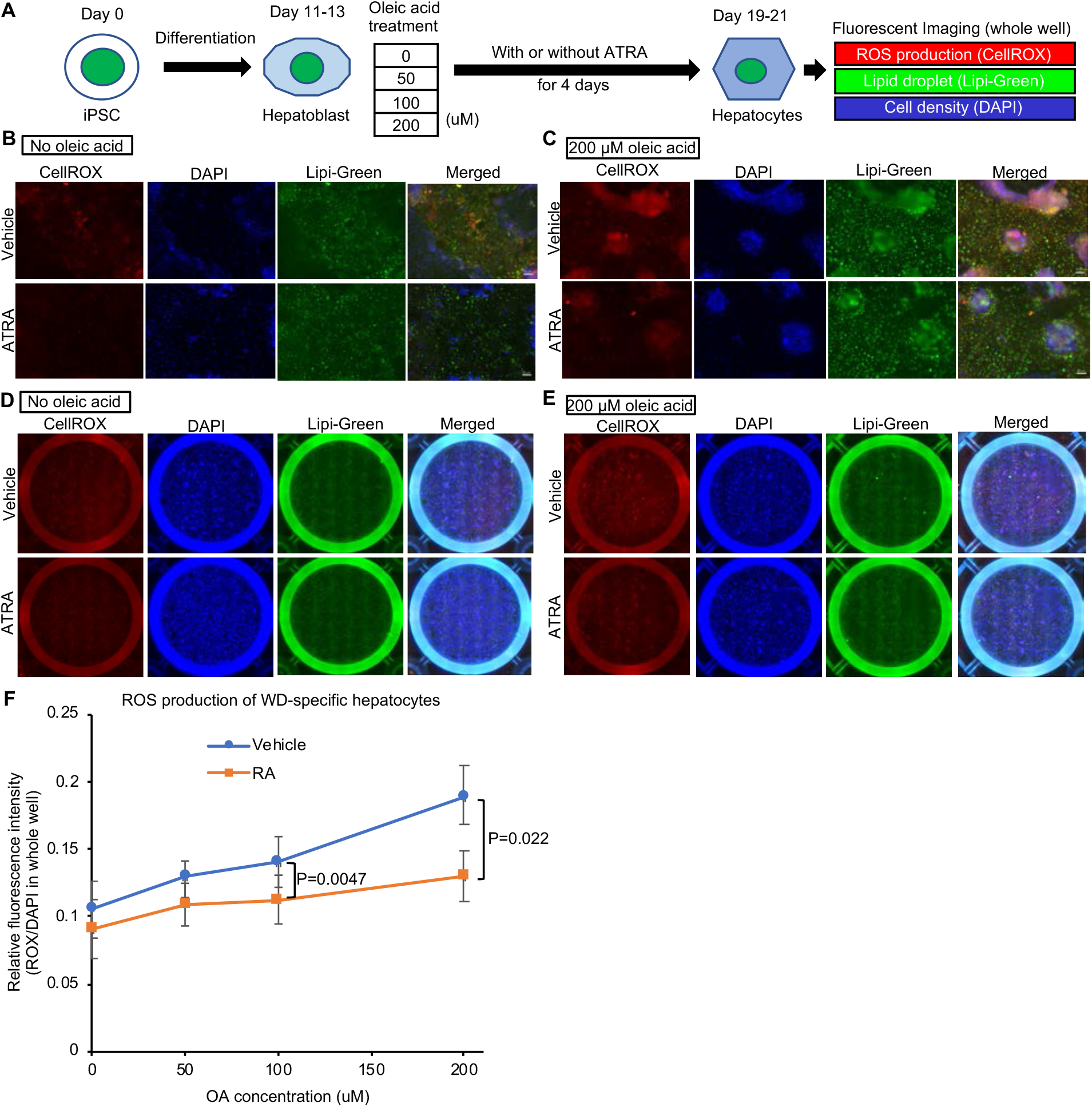
ATRA decreases oleic acid-induced ROS production in WD-iPSC-derived hepatocytes. (A) Schema of a cellular assay of ROS production and lipid accumulation in WD-iPSC-derived hepatocytes treated with different concentrations of oleic acid with or without ATRA. (B-C) Magnified lipid fluorescent images of CellROX, DAPI, and Lipi-Green in HPS0045-derived hepatocytes on differentiation day 19 after oleic acid treatment at different concentrations for 8 days. Scale bars, 50 µm. (D-E) Tiled whole-well fluorescent images of CellROX Deep Red, DAPI, and Lipi-Green in HPS0045-derived hepatocytes after oleic acid treatment at different concentrations for 8 days with or without ATRA for 4 days. (F) Quantified ROS production at different concentrations of oleic acid with or without ATRA in four WD-patient-derived hepatocytes (HPS0045, HPS0049, HPS0053, and HPS2807). Data are shown as mean ± SEM (n=4). P-values were determined by two-tailed Student’s *t*-tests for paired samples from each cell line.

## Discussion

In this study, we found that ATRA and some other clinically-approved retinoids rescued the decreased expression and secretion of Cp in WD-specific and ATP7B-deficient hepatocytes. Furthermore, ATRA alleviated ROS production in WD-specific hepatocytes treated with higher concentrations of oleic acid. Because Cp reduction and liver steatosis are the initial symptoms of WD, our findings suggested that retinoids could be new therapeutic drugs to prevent aggravation of WD. This therapeutic approach can be supported by previous studies using genetically modified animals. Loss of RA signaling in mouse liver resulted in steatohepatitis and liver tumors, and feeding on a high RA diet rescued hepatic abnormalities and prevented liver tumors (Yanagitani et al., 2004). RA also exhibited suppressive effects on iron-induced oxidative stress in the same transgenic mouse (Tsuchiya et al., 2009). Also, ATRA effectively improves liver steatosis in a rabbit model of high fat (Zarei et al., 2020). In Atp7b(-/-) mice, the activity of LXR (liver X receptor)/RXR pathway decreases (Wooton-Kee et al., 2020), and the activation of these pathways or the feeding of zinc-enriched foods could ameliorate liver damages (Buiakova et al., 1999; Hamilton et al., 2016; Wooton-Kee et al., 2015). Together with our transcriptome analysis on WD-specific hepatocytes, which showed decreased RA signaling pathways and increased lipid metabolism, our findings represent pieces of evidence on the efficacy of retinoids to prevent hepatic steatosis caused by WD using patient-derived hepatocyte models.

We have generated iPSC lines from 4 WD patients, which carry compound heterozygous mutations, as well as ATP7B-KO iPSC lines. We have also corrected the mutation in an existed iPSC line carrying R778L homozygous mutation (Zhang et al., 2011). Some of the mutations that we report here were not previously reported, nor the complicated compound effects of these combinations of the mutations. Thus their genotype-phenotype correlations were still under investigation. Previous studies using WD-specific iPSCs, which used only one patient line each, just focused on the expression, localization, and functionality of ATP7B and the cellular toxicity might cause to show different results because heterogeneous expression levels of ATP7B protein among these WD-specific hepatocytes as we have shown in this study. Our findings demonstrated that the common pathological mechanisms of WD can be obtained from the integrative approaches using various genotypes in *ATP7B* gene.

We found that Cp expression and secretion are robustly decreased in WD-specific hepatocytes. Since low Cp concentration in the plasma is a general clinical feature of WD, our findings represent a successful recapitulation of WD phenotypes *in vitro*. Interestingly, not only Cp secretion level but also mRNA expression level of Cp was down-regulated in WD-specific hepatocytes. These results cannot be explained by the common concept that apo-ceruloplasmin has weak protein stability compared to holo-ceruloplasmin, in which copper ion is delivered by ATP7B. Indeed, a previous study reported that Cp mRNA expression in the liver sample of WD patients was decreased (Czaja et al., 1987). Decreased Cp mRNA must be regulated by other mechanisms in WD-specific hepatocytes. Also, Cp expression was reported to be decreased by an mRNA decay mechanism in response to the intracellular oxidative stress (Tapryal et al., 2009). Thus, decreased Cp mRNA expression in WD-specific hepatocytes might be caused by the abnormal oxidative stress and/or nuclear receptor signaling pathways.

We have developed a Cp secretion-based drug screening assay and a ROS detection assay in response to high oleic acid concentration using human iPSC-derived hepatocytes. Our strategy may represent a faithful recapitulation of initial cellular features of WD-specific hepatocytes and thus provide effective platforms to develop potential therapeutics for hepatic steatosis in WD and other fatty liver diseases as well as to examine the molecular pathogenesis of WD. Because the detection methods are clear and easy, these assays should be further sophisticated to allow high throughput screening systems to identify the ideal therapeutic drugs in the future.

### Limitation of study

Although we have demonstrated that retinoids rescued ceruloplasmin secretion and alleviate oxidative stress in Wilson’s disease-specific hepatocytes for the first time, animal tests and preclinical evaluation will be required to proceed to clinical trials. Also, although we have used iPSC lines from enough numbers of WD patients for statistical tests to draw the general conclusions for the first time, there might be outliers due to the variation of WD patients with different types of mutations. Last, the molecular mechanisms of how the ATP7B deficiency leads to abnormal lipid mechanisms remain elusive. It is interesting to investigate the relationships of copper and lipid metabolisms in the future.

## Acknowledgments

We thank Drs. Miguel A Esteban and Duanqing Pei (Chinese Academy of Sciences) for providing us with the ATP7B R778L homo iPSC line and Dr. Bruce Conklin for providing us with WTC11 iPSC line. We would like to express our sincere gratitude to all our coworkers and collaborators, to Kumiko Omori for their administrative support.

This research was supported by the Japan Agency for Medical Research and Development through its research grant “Core Center for iPS Cell Research, Research Center Network for Realization of Regenerative Medicine”. This research was supported in part by the grants from a JSPS KAKENHI Grant-in-Aid for Young Scientists (A) (17H05063) to Y.H., Grant-in-Aid for Early-Career Scientists (18K15054) to G.T., Grant-in-Aid for Scientific Research (B) (20H03442) to Y.M., Grant-in-Aid for Challenging Research (Pioneering) (20K21409) to Y.M., Grants for iPS cell research from AMED (12103610 and 17935400) to M.K.S., Interstellar Initiative from AMED (20jm0610032h0001) to Y.M., Takeda Science Foundation to Y.H. and Y.M., Uehara Memorial Foundation to Y.H., The Tokyo Biochemical Research Foundation to Y.H. and Y.M., Sumitomo Foundation Grant For Basic Science Research Projects to Y.M., Ichiro Kanehara Foundation Grant to Y.M., and Naito Foundation Research Grant to Y.M.

## Author contributions

Conceptualization, Y.H. and Y.M; Methodology, D.S., G.T., Y.W.Z., M.M.T., J.L., M.T., Y.A., Y.H., Y.M., and Y.H.; Investigation, D.S., G.T., Y.W.Z., M.M.T., J.L., M.T., Y.A., Y.H., Y.M., and Y.H.; Resources D.S., G.T., N.M., T.F., M.N., T.N., M.K.S., Y.N., Y.M, and Y. H; Writing – Original Draft, D.S., G.T., Y.M. and Y.H.; Writing – Review & Editing, D.S., G.T., Y.M. and Y.H.; Visualization, D.S., G.T., Y.M. and Y.H.; Supervision, Y.W.Z., T.O., Y.M. and Y.H.; Project Administration, T.N., M.K.S., N.Y., T.O., Y.M. and Y.H, Funding Acquisition, Y.M. and Y.H.

